# SKIOME Project: a curated collection of skin microbiome datasets enriched with study-related metadata

**DOI:** 10.1101/2021.08.17.456635

**Authors:** G. Agostinetto, D. Bozzi, D. Porro, M. Casiraghi, M. Labra, A. Bruno

## Abstract

Large amounts of data from microbiome-related studies have been (and are currently being) deposited on international public databases. These datasets represent a valuable resource for the microbiome research community and could serve future researchers interested in integrating multiple datasets into powerful meta-analyses. However, this huge amount of data lacks harmonization and is far from being completely exploited in its full potential to build a foundation that places microbiome research at the nexus of many subdisciplines within and beyond biology. Thus, urges the need for data accessibility and reusability, according to FAIR (Findable, Accessible, Interoperable, and Reusable) principles, as supported by National Microbiome Data Collaborative and FAIR Microbiome.

To tackle the challenge of accelerating discovery and advances in skin microbiome research, we collected, integrated and organized existing microbiome data resources from human skin 16S rRNA amplicon sequencing experiments. We generated a comprehensive collection of datasets, enriched in metadata, and organized this information into data frames ready to be integrated into microbiome research projects and advanced post-processing analysis, such as data science applications (e.g. machine learning). Furthermore, we have created a data retrieval and curation framework built on three different stages to maximize the retrieval of datasets and metadata associated with them. Lastly, we highlighted some caveats regarding metadata retrieval and suggested ways to improve future metadata submissions.

Overall, our work resulted in a curated skin microbiome datasets collection accompanied by a state-of-the-art analysis of the last 10 years of the skin microbiome field.

## Introduction

Directly in contact with the environment, the skin microbiome is a tangled and dynamic ecosystem that interacts with both the host and its surroundings (*1*). It is characterized by diverse ecological niches, where the microbiota, the host skin cells and the host immune system are involved in the maintenance of skin health. In the last decade, numerous studies have investigated the composition of the human skin microbiome under very different conditions (*2–4*).

The advent of high-throughput DNA sequencing (HTS) technologies has revolutionized numerous research fields, and the study of the human microbiome was no exception. Following the introduction of HTS technologies, the number of studies investigating the human microbiome has increased, expanding our knowledge about its implications for human health. In particular, it was demonstrated its pivotal linkage with diet and age (*5,6*) and specific microbiome patterns were shown to relate to the body region sampled (*7,8*). Geography and ethnicity have also been shown to affect the skin microbiome (*9*) and numerous diseases have been associated with an altered microbial state (*10*), as in the cases of atopic dermatitis (*11*) and psoriasis (*12*).

Since their adoption, the new sequencing strategies have been getting cheaper and cheaper, becoming available for researchers and companies on a global scale. In recent years, large amounts of data have been deposited in public databases and more is going to be produced in the near future, as the number of sequencing experiments is exponentially growing.

There are three major databases used to store nucleotide sequence data: the NCBI’s Sequence Read Archive (SRA) (*13*), the EBI’s European Nucleotide Archive (ENA) (*14*), and the DDBJ Sequence Read Archive (DRA) (*15*). These three databases are brought together by the International Nucleotide Sequence Database Collaboration (INSDC) and are constantly synchronized to share their data (*16*). The publicly available datasets deposited in these databases represent a valuable resource for the microbiome research community. Public available data can be now accessed and downloaded to be re-analysed or integrated to perform meta-analysis studies (*17–19*).

As a consequence, in the last few years, we are facing an increasing adoption of novel large-scale data science approaches to address challenges in microbiome science (*20*). For example, machine learning strategies can be applied to perform powerful prediction tasks on metagenomics data (e.g. disease-prediction based on microbiome composition). However, these strategies require a large amount of data to train and test models, making the integration and harmonization of multiple datasets a necessary step (*21,22*). In this way, the availability of large-scale sequencing data can enable microbiology researchers to ask new questions and develop new strategies to study the human-associated microbial communities (*23,24*).

However, this huge amount of microbiome data still lacks harmonization and is far from being completely exploited to its full potential. Guidelines have been proposed and tools have been developed to promote the standardization of sample processing, sequencing and data analysis across the microbiome field (*25–32*) but achieving global standardization is not an easy task. Initiatives such as the Human Microbiome (*33*) and the Earth Microbiome Projects (*34*) have favored the development of standardized procedures. In addition, important field-specific databases were created, such as the Human Oral Microbiome Database (*35*) or the GMrepo, a database of curated and consistently annotated human gut metagenomes (*36*).

Several research groups have been proposing different sources of microbiome data: initiatives like the Human Microbiome and the Integrative Microbiome Projects (*37,38*), MicrobiomeDB (*39*), HumanMetagenomeDB (*40*), curatedMetagenomicData (*41*), the ML Repo (*42*), QIITA portal (*43*), or the MG-RAST portal (*44*) suggested both data management infrastructures and frameworks to guarantee data accessibility and reuse.

Despite the contribution of groups involved in this field, the lack of metadata and the presence of datasets with missing or inconsistent information can reduce the interpretability of the data generated, influencing the understanding of microbial dynamics and ecological patterns (*23,24,45*). Inconsistency and uncontrolled metadata filling were demonstrated by Gonçalves and Musen (*46*), revealing the necessity of standardized metadata compilation (*47*).

FAIR (Findable, Accessible, Interoperable, and Reusable) principles are supported within the National Microbiome Data Collaborative and FAIR Microbiome community (https://www.go-fair.org/implementation-networks/overview/fair-microbiome) (*23,45*) to promote data discovery and reuse in the microbiome field, and allow for broader dissemination of knowledge and compliance for both humans and machines.

Thus, making microbiome data and metadata accessible is a key aspect to guarantee a concrete opportunity to perform meta-analyses and data reuse (*42,48,49*). In this context, well-curated and FAIR microbiome datasets are now a necessity to explore microbiome patterns, apply data science techniques and promote data reusability (*50,51*).

In order to help researchers interested in performing meta-analyses with human skin microbiome data and exploring the context-specific information related to potentially useful datasets, we focused our work on published human skin microbiome datasets, creating a curated skin microbiome collection accompanied by a state-of-the-art analysis of the last 10 years of the skin microbiome field.

In particular, during the last decade, most of the studies have relied on amplicon sequencing approaches, where different regions of the 16S rRNA gene are amplified and sequenced to identify the microbial taxa present in a sample (*52,53*). For this reason, we built a comprehensive human skin microbiome collection enriched with detailed metadata information, focusing on existing 16S rRNA amplicon-sequencing microbiome datasets from the human skin biome.

To achieve our goal, we first collected datasets from the INSDC, which store the majority of the publicly available nucleotide sequencing datasets together with their associated metadata (*16*). As the availability of these metadata and the possibility of recovering them is crucial for ensuring the reusability of the available datasets (*46*), we dedicated special attention to maximize the amount of metadata information that can be recovered. To do so, we combined different metadata retrieval approaches enriched with a manual curation step. Then, we generated explorable data frames at different curation levels containing all the retrieved datasets together with the associated metadata. Further, we highlighted some of the shortcomings of the current approaches for data and metadata retrieval and we called attention to some of the issues that currently afflict the re-usability of the deposited data. Overall, the output of our work constitutes a valuable resource for researchers interested in performing meta-analyses with human skin microbiome data, who can explore our collection to find a list of datasets that can be integrated to answer old and new biological questions.

## Materials and methods

### Metadata retrieval and manual curation procedures

To obtain a comprehensive list of skin microbiome studies derived from amplicon approaches with the associated metadata, we built a three-step framework (Fig. 1) based on:

- Step 1: dataset retrieval from INSDC;
- Step 2: metadata retrieval and enrichment;
- Step 3: output curation with the removal of redundant and spurious information.

**Figure 1.**
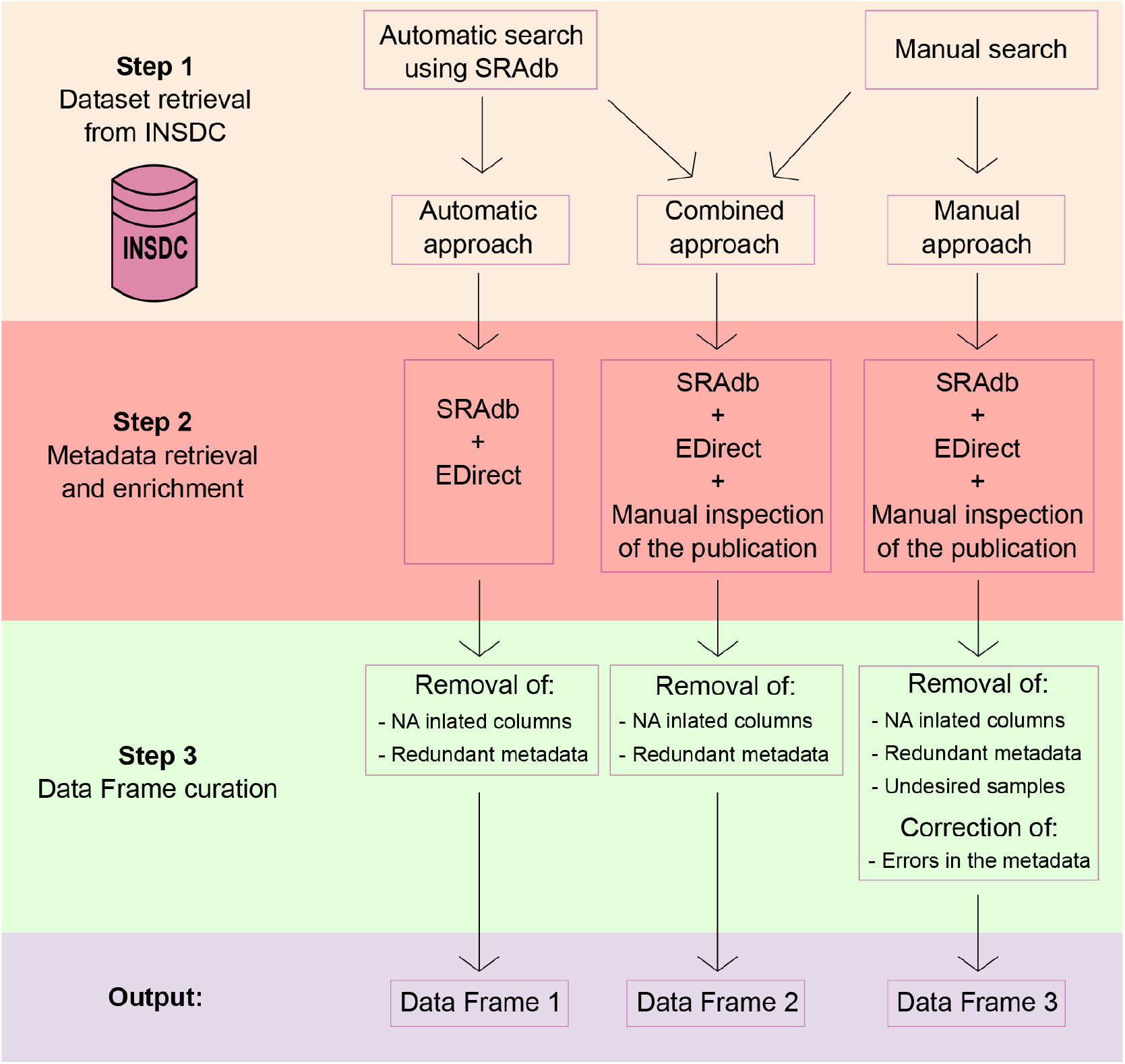
Schematic representation of the three-step framework adopted in the study to collect datasets and metadata and generate three differently curated data frames.

In the sections below, all the steps are described together with the methods and strategies used.

### Step 1: dataset retrieval from INSDC

To generate a comprehensive list of datasets of human skin microbiome derived from 16S rRNA amplicon sequencing available on the INSDC public databases, we decided to rely on two different approaches: i) an automatic search, which allows querying the INSDC databases automatically using keywords and ii) a manual approach on the SRA and ENA portals.

The automatic search of the datasets was performed with the R package “SRAdb” (*54*). SRAdb relies on a SRAdb SQLite database, a regularly updated database of metadata associated with the raw reads deposited on SRA and its interconnected databases (ENA, DRA). The SRAdb database (up to 36 Gb) was downloaded and stored locally on the 17^th^ of June, 2021. We performed a full-text search with the following query: “human skin microbiome OR human skin microbiota OR human skin metagenome”.

For the manual approach, instead, we performed a search on the NCBI’s SRA and EBI’s ENA databases with the following criteria: datasets coming from 16S rRNA amplicon sequencing, containing only human skin samples that were deposited from 2012 onwards and that presented an associated publication.

### Step 2: metadata retrieval and enrichment

An enrichment step was performed on both automatic and manual outputs in order to recover the largest amount of metadata associated with the datasets previously found. For this step, we integrated three different strategies: i) SRAdb was used to collect all the possible information from the retrieved list of studies and samples; ii) for some run-associated metadata that could not be retrieved with SRAdb, we used the Entrez Direct (EDirect) tool (*55*); iii) for the list of manually recovered studies, we collected study-specific metadata from the associated publication, including information that cannot be found on the INSDC databases. We focused our attention on the sample origin, the laboratory and bioinformatics strategies and the data related to the context in which the studies were performed. In particular, we retrieved study-specific information related to the collection method used, the 16S rRNA gene hypervariable region sequenced, the clustering method used (OTUs, ASVs/RSVs), the number of recovered units/variants reported in the study, the database used for taxonomic assignment and its version, the disease condition investigated (if any), the location of the sampling, the presence of a MGnify analysis (*56*), the DOI and the year and journal of publication.

In addition, a bibliometric analysis of published papers related to the datasets retrieved was performed. Research areas and categories from the Web of Science (WoS) collection and Elsevier’s Scopus classifications were added to each publication. Notably, since Scopus reported multiple subject areas for each publication, we included multiple columns in the data frame to keep all the information. We further generated a column categorizing a scientific journal as a medicine-related journal (Medicine_Journal) or not depending on the presence of ‘Medicine’ among the Scopus subject areas. Lastly, an additional column containing any useful notes related to the study was added.

A comprehensive list of the manually curated metadata with description is available in Supplementary File 1., also available in our Github repository (https://github.com/giuliaago/SKIOMEMetadataRetrieval).

### Step 3: outputs curation and metadata correction

Once all the information was stored into three data frames that differed in the way the datasets and the metadata were retrieved, we proceeded to reorganize them by removing redundant metadata and NA-inflated columns. For the smallest and most refined data frame, we further inspected the data frame rows to remove undesired samples and to correct wrongly assigned metadata. In detail, we removed samples that were not obtained from amplicon sequencing and corrected metadata by double-checking with the related publications.

### Script and data availability

For all the steps of datasets and metadata retrieval, a list of studies and associated metadata were kept (Dataframe 1, Dataframe 2 and Dataframe 3). All the outputs will be available in our Github repository (https://github.com/giuliaago/SKIOMEMetadataRetrieval), accompanied by the scripts used for the retrieval framework. In particular, scripts describe the use of SRAdb, Edirect tool, the entire R pipeline to obtain the final outputs and codes for plot creation and data frame exploration.

## Results

Following the three steps presented in the Methods section (dataset retrieval from INSCD; metadata retrieval and enrichment; data frame curation), we first tested two approaches to retrieve datasets of the human skin microbiome from the INSCD databases (Step 1): a manual search of the datasets and an automatic search with SRAdb (*54*). We then collected metadata information for the retrieved datasets (Step 2) using three different approaches: automatic search with SRAdb (*54*), EDirect (*55*) and a manual search from the associated publication for the manually retrieved studies. In this way we obtained three data frames:

- Data Frame 1, containing only datasets retrieved with SRAdb and metadata collected automatically with SRAdb and EDirect;
- Data Frame 2, containing all the datasets identified with both the strategies (manual and automatic) together with all the metadata that could be recovered with SRAdb, EDirect and manual inspection of the publication;
- Data Frame 3, a subset of Data Frame 2, containing only the manually retrieved datasets together with all the metadata that could be recovered both manually and automatically with SRAdb and EDirect.

Data Frame 2 and Data Frame 3 both contain 61 metadata columns (from manual and automatic metadata search), while Data Frame 1 only contains 37 metadata columns obtained from the automatic search. All three data frames were curated to remove redundant columns and NA-inflated columns (Step 3). Among the redundant metadata, we observe columns containing the IDs of Run, Experiment, Submission, Sample/BioSample and Study/BioProject. Other metadata recovered by both methods were the spots, the bases, the library strategy, the sequencing platform used and the Taxon ID. Data Frame 3 was further curated to remove undesired samples coming from whole-genome sequencing experiments and to correct wrongly assigned metadata.

The following sections will show the results, starting from a comparison between the data collection approaches used and then moving to describe the state-of-the-art of metadata related to the submission process and the metadata obtained from our manual curation step, in particular regarding the bioinformatic strategies used and the skin data characteristics retrieved directly from the published studies.

### Comparison of datasets collection approaches and metadata retrieval

The automatic search with SRAdb recovered a total number of 97,182 samples from 203 studies (Data Frame 1) with 8,492 samples that were uploaded before 2012. The manual search, instead, recovered a total of 21,958 samples from 68 studies (Data Frame 3) starting from 2012.

We compared the ability of the two approaches in identifying the desired datasets. Notably, the automatic search failed to identify 47 studies that were recovered by the manual search, indicating that SRAdb does not perform an exhaustive search of the available datasets. The automatic search identified 182 studies not found by the manual search. Based on these observations we generated a data frame (Data Frame 2) that comprised both automatically retrieved and manually identified studies. This data frame contains 108,207 rows (samples) coming from 250 different studies and a total of 61 columns containing the metadata.

The metadata associated with the datasets can be differentiated into three major categories: i) metadata related to dataset submission (obtained by the automatic search), ii) metadata associated with the laboratory procedures and bioinformatic pipelines (obtained by automatic and manual searches) and iii) manually collected context metadata describing other relevant aspects of the study (e.g. disease/condition investigated or sample origin).

The automatic search for metadata with SRAdb and EDirect was performed for all the datasets, both manually and automatically retrieved, to collect metadata related to dataset submission (i). After the curation step, we conserved a total of 37 metadata columns that were included in all three data frames.

These 37 columns contain information related to:

- the study with BioProject, Study_ID, Study_description and Study_abstract;
- the submission and its date with the Year_of_release, Release_Date and Load_Date;
- the experiment with the Library Strategy used (Library_Strategy), specification on if it was performed a pair-end or a single-end sequencing (Library_Layout) and the library Insert size (Insert_Size);
- the sequencing platform and the model used (Platform, Model);
- the run with the average sequence length (AvgLength), the spots, the bases, the size of the file (Size_MB) and the path for the download (Download_path);
- the experiment title (Experiment_title);
- a description of its design (Design_description);
- the name of the library (Library_name) and attributes of the experiment (Experiment_attribute);
- the sample with BioSample, Sample_ID, Sample_alias, Sex, Body_Site, Description and Sample_attribute and
- the associated Taxonomic ID with the scientific name (TaxID, Scientific_Name).

A comprehensive description of all the 37 metadata is available in Supplementary File 1.

In Data Frame 2 and 3 we also included 23 additional columns that contain metadata not available on INSDC and obtained from the manual inspection of the publication. These metadata were recovered only for the manually retrieved datasets and contained information on the laboratory procedures and bioinformatic pipelines (ii) together with other relevant metadata describing the context of the study (iii).

In the next sections, all the categories of metadata and their distribution are outlined. A full description of the metadata included in the data frames is given in Supplementary File 1, also available in our Github repository (https://github.com/giuliaago/SKIOMEMetadataRetrieval).

### Distribution of metadata related to dataset submission and library preparation

By comparing the distribution of the number of datasets released over the years among the three different data frames (Fig. 2.b), we observed that Data Frame 1 showed a peak in 2015 when 17,551 datasets were released. Differently, Data Frame 2 showed a peak in 2017 with 19,041 datasets released during that year. For Data Frame 3, we observed two peaks: one in 2013 with 4,841 datasets released and one in 2017 with 7,293 datasets released. However, if we look at the number of studies, the peak was reached in 2019 with 16 studies investigating the human skin microbiome (Fig. 2a).

**Figure 2.**
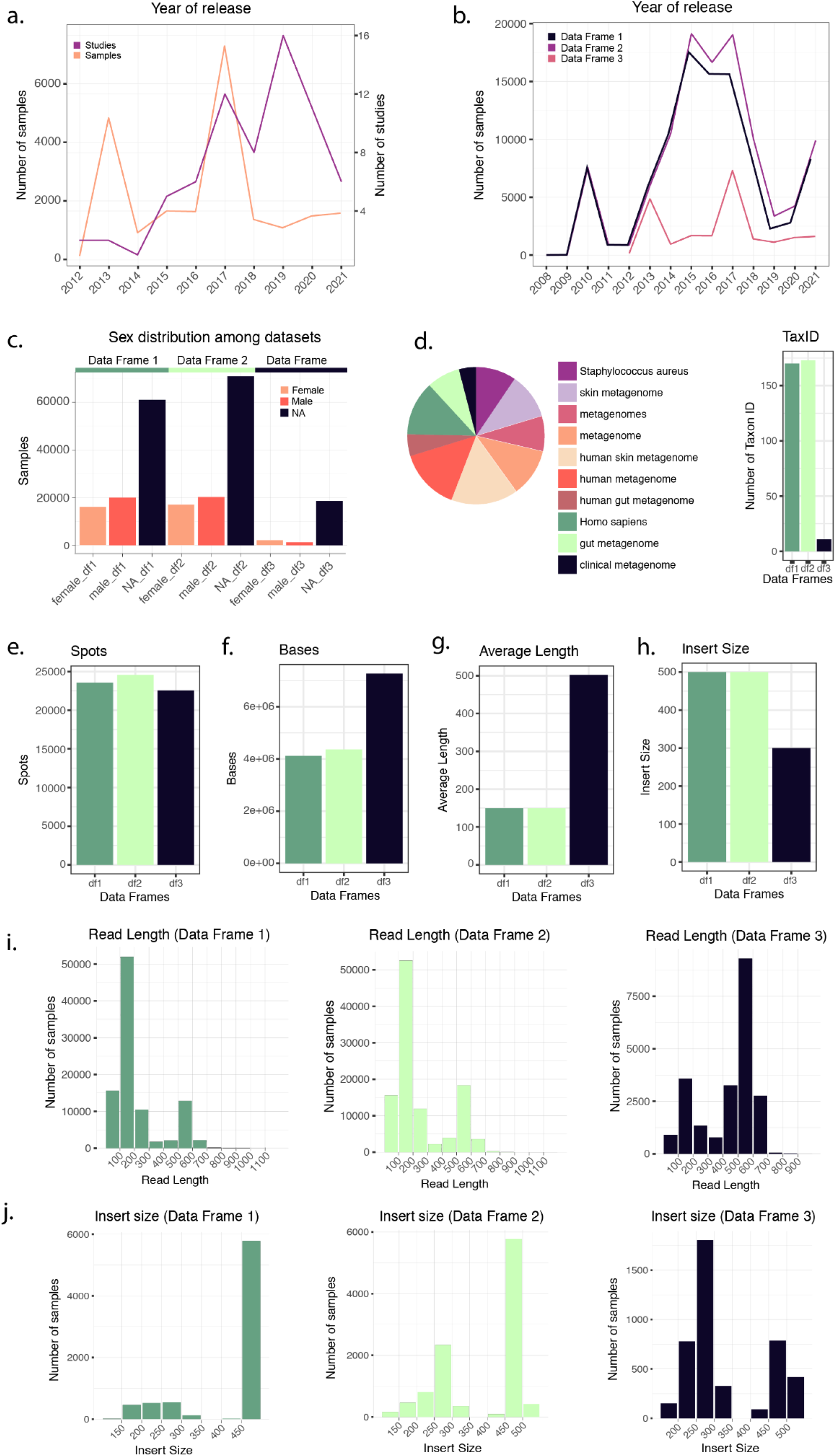
**a**) Number of studies and samples from Data Frame 3 released every year starting from 2012. **b**) Comparison of the number of samples released each year for the three Data Frames (Data Frame 1 in blue, Data Frame 2 in black and Data Frame 3 in red). Data Frames 1 and 2 contain samples starting from 2008, while Data Frame 3 only from 2012. **c**) Distribution of the variable “sex” in the three Data Frames. In all three cases, the majority of the samples don’t have such information reported. **d**) The number of Taxon ID/Scientific names used in the three Data Frames (barplot) and relative abundance (as a logarithm) of the Taxon ID/Scientific names used for the samples in Data Frame 3 (pie chart). **e-h**) Comparison of the median number of spots (**e**), bases (**f**), reads average length (**g**) and insert size (**h**) in the three Data Frames. **i**) Read length distribution in the three Data Frames. **j**) Distribution of the insert size in the three Data Frames.

After removing datasets with a value equal to zero for the following metadata, we calculated the median number of spots (sequencing clusters that generated sequence), bases (nucleotides), average read length and insert size (size of the amplicon without sequencing adapters) for Data Frames 1, 2, and 3. The median number of spots were respectively 23,590, 24,564, 22,560.5 (Fig. 2e), while the median number of bases were 4,114,610, 4,364,032 and 7,270,396 (Fig. 2f). The mean of the datasets’ average read length in Data Frame 1 is 227.0235 bp, while for Data Frame 2 is 254.0603 bp and for Data Frame 3 is 440.2783 bp. The median values are 150 bp for Data Frame 1 and 2, and 502 bp for Data Frame 3 (Fig. 2g and Fig. 2i). The median insert size is 500 in Data Frame 1 and 2 and 300 in Data Frame 3 (Fig. 2h and Fig. 2i). Mean values are 455.5963, 440.2783 and 349.0783, respectively.

Information about the sex of the individuals can be collected for 36,231 out of 97,182 samples in Data Frame 1 (20,011 females; 16,220 males), 37,340 out of 108,207 samples in Data Frame 2 (20,234 females; 17,106 males), and 3,461 out of 21,958 samples in Data Frame 3 (1,276 females; 2,185 males).

We recognized 66 different descriptions (more or less accurate), defining the sampled region of the body. However, metadata on the body site is absent in most of the datasets. In detail, a total of 42,489 empty metadata information were found for Data Frame 1, 52,972 for Data Frame 2 and 18,061 for Data Frame 3.

In our data frames, we have observed the use of different Taxon IDs to describe the samples. Data Frame 3, which contains only samples of human skin microbiome, presents 11 different taxon IDs, which correspond to the following scientific names: “human skin metagenome”, “*Homo sapiens*”, “metagenome”, “metagenomes”, “human metagenome”, “skin metagenome”, “*Staphylococcus aureus*”, “clinical metagenome”, “gut metagenome”, “human gut metagenome” and “bacterium”. The number of Taxon IDs increases in the other two data frames so that in Data Frame 2 we observe 173 different Taxon IDs.

### Methodological pipeline insights and context-metadata of skin microbiome datasets

For the 68 manually retrieved studies we further collected other metadata from the associated publications. Based on these manually collected metadata, we observed that most of the studies had used swabs to collect samples (53 studies; 19,928 samples), with only a few relying on other methods like biopsies (5 studies; 257 samples), scrubs buffer washes (1 study; 1,358 samples) or a combination of swabs and other methods (7 studies; 311 samples).

Considering the marker gene used, the most commonly sequenced hypervariable regions of the 16S rRNA gene have been the V1-V3 (6,176 samples), followed by the V4 (*5,694*) (Fig. 3a). However, if we consider the number of studies, we observed that most of them relied on the V1-V3 (24 studies) and V3-V4 (21 studies) regions (Fig. 3a). The Illumina sequencing platforms were the most used (88,295 samples in Data Frame 2), particularly the Illumina Miseq platform (49,297 samples in Data Frame 2), followed by Roche 454 platform (19,777 samples in Data Frame 2). A total of 11,412 samples have no specific platform model assigned (Fig. 3c).

**Figure 3.**
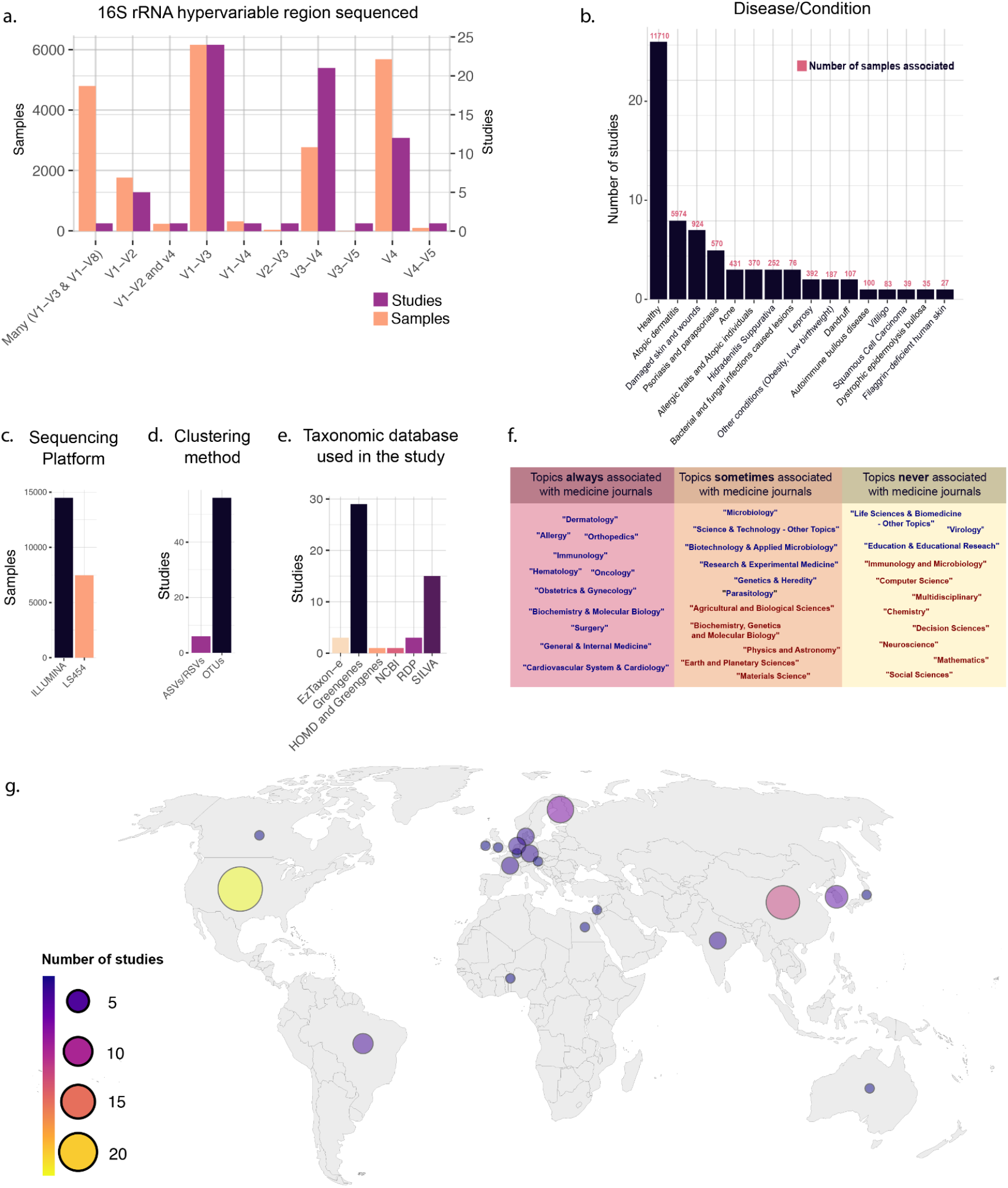
**a**) Number of samples (pink) and studies (purple) that used specific 16S rRNA hypervariable regions in Data Frame 3. **b**) The number of studies and samples for each disease/condition investigated in Data Frame 3. **c-e**) Frequency of use of the different sequencing platforms (**c**), clustering methods (**d**) and taxonomic databases (**e**) in Data Frame 3. **f**) Table showing the Web Of Science research areas (blue) and Scopus Research Subjects (red) that described the scientific journals in which the studies of Data Frame 3 have been published. The research areas/subjects are divided into three boxes depending on how often they were associated with the Scopus research subject “Medicine”. Going from left to right are shown the research areas/subjects that were always (left), sometimes (center) and never (right) associated with the Scopus research subject “Medicine”. **g**) Geographical distribution of the studies included in Data Frame 3.

Regarding the bioinformatic pipeline used, most of the manually inspected studies have clustered reads into Operational Taxonomic Units (OTUs) (56 studies), only a few (6 studies) relied on Amplicon Sequence Variants (ASVs) or Ribosomal (*35*) Sequence Variants (RSVs). For 6 studies this information was not reported in the article methods (Fig. 3d).

Taxonomy assignment was mainly performed with Greengenes database (*57*) (29 studies), followed by SILVA (*58*) (15 studies). Other works relied on different databases, including RDP (*59*) (3 studies), EzTaxon-e (*60*) (3 studies), NCBI (1 study), and HOMD (1 study). Strikingly, many studies did not report this information in the articles’ method section (16 studies) (Fig. 3e).

Our analysis also comprehended a detailed inspection of skin and disease conditions related to the microbiome analysis. Among our list, we identify 42 studies investigating 26 different diseases/conditions of the skin (Fig. 3b). The most commonly investigated disease in our curated dataset is atopic dermatitis (8 studies), followed by psoriasis and parapsoriasis (5 studies), while 7 studies investigated skin injuries of different kinds. Among the other diseases/conditions investigated, we observed acne (3 studies), skin pathogenic infections, such as bacterial, fungal and parasitic infection (3 studies), allergic traits and atopic individuals (3 studies), dandruff (2 studies), leprosy (2 studies), hidradenitis suppurativa condition (2 studies), autoimmune bullous disease (1 study), dystrophic epidermolysis bullosa (1 study), vitiligo (1 study), squamous cell carcinoma (1 study), filaggrin-deficient human skin (1 study), and other conditions such as obesity and low birth weight (2 studies). Overall, 26 studies collected samples from healthy human skin (in Data frame 3, column 43 ‘disease/condition’).

Looking at the geographic distribution of the studies, we observed that most of them were conducted in the USA (22 studies), followed by European countries (19 studies) and China (11 studies). Other countries that featured more than one study were South Korea (4 studies), Brazil (3 studies) and India (2 studies) (in Data frame 3, column 44 ‘Location’) (Fig. 3.g).

Finally, the 68 manually retrieved studies were published in 40 different scientific journals from 17 different WOS research areas. According to Scopus classification, 36 studies were published in medicine-related scientific journals (Research Subject = Medicine). Figure 3f shows how often specific WOS Research areas and Scopus Research Subjects are associated with the Scopus research subject “Medicine” in the present dataset.

## Discussion

In this section we discuss the results obtained from our work, in particular focusing the attention on three main aspects: i) outcomes related to dataset collection, ii) caveats related to metadata retrieval and data reuse and, finally, iii) the importance of having a curated collection of a microbiome dataset for advancing the microbiome research field through data-driven approaches and powerful meta-analysis.

### Skin microbiome data retrieval: dataset collection is not an easy task

The INSDCs databases are the source of an enormous amount of publicly available datasets which can be accessed and downloaded to perform powerful meta-analyses (16). The field of microbiome research can greatly benefit from the availability of this large amount of data (23). However, the reusability of a dataset strictly depends on the possibility of retrieving it and on the amount of information (metadata) deposited by the authors at the time of submission (*46,61*).

If the number of datasets available is limited (such as for poorly studied environments), a manual search will consent to gather all the studies available in a relatively fast way. However, for well-studied environments, the number of datasets can be very large and it becomes more convenient to rely on automatic approaches (*62*). The automatic approach allows for a fast and comprehensive search of datasets of interest, but at the same time, it lacks a curation step that validates the recovered datasets. Moreover, the automatic search does not permit the retrieval of important information that was not deposited in the INSDC databases together with the raw data. Conversely, the manual search is more accurate and allows a researcher to retrieve a well-validated list of studies together with other information by inspecting the associated publication. Its drawbacks are that it is time-consuming and presumably less comprehensive than the automatic search. Moreover, it does not consent to retrieve sample-specific information.

Our results showed that the automatic search did find a greater number of datasets than the manual (97,182 samples from 203 studies vs 21,958 samples from 68 studies). Many can be the reasons that explain this difference. First, the automatic search tends to be more exhaustive than a manual one if the number of available datasets is large. Second, the list of studies is not inspected to remove undesired studies that do not match some of the desired criteria but might be retrieved by the searching tool. Third, the manual search was limited to the dataset deposited in the last 10 years, starting from 2012, while the automatic search recovered studies starting from 2008. Indeed, 8,492 samples found by the automatic search were uploaded before 2012. Despite these observations, neither the manual nor the automatic search with SRAdb, were capable of recovering all the studies, highlighting the importance of combining the two approaches.

Together, our results indicated that SRAdb was not exhaustive in its search, and to maximize the number of datasets retrieved, a combination of manual and automated approaches might represent the optimal strategy. We observe that the larger the number of available datasets, the less feasible an extensive manual search, favoring an automated approach for the dataset retrieval step. Conversely, for topics with a particularly small number of datasets available, the manual search still remains the most accurate way of recovering them.

### Caveats of metadata retrieval and data reuse

Depending on the topic, a researcher interested in performing a meta-analysis can decide to rely on different approaches to retrieve metadata associated with the datasets of interest, both directly through the INSDC data portal (*16*) or with specific tools (*54,55,63*). In this work, we decided to combine three approaches, based on SRAdb (*54*), Entrez (*55*) plus a manual search from the publication, with the aim of generating a comprehensive data frame containing all the datasets from the human skin microbiome amplicon sequencing available on INSDC databases. As for the search of the datasets, also for metadata retrieval, we observed that the combination of automatic and manual approaches is capable of gathering a larger amount of information than the two approaches alone.

However, while with a manual search it is possible to recover much information related to a dataset if a publication is available, this approach is not feasible if the number of datasets is high (*62*). Moreover, sample-specific information for large datasets can only be collected using automatic approaches, making an automatic search a necessity.

Automatic approaches of metadata retrieval (such as those used in this study) collect the metadata deposited on the INSDC databases. As such, they are capable of accessing only the metadata that were made available by the researchers during the data submission. Failing in accessing specific metadata can affect the re-usability of a given dataset, highlighting the importance of proper and extensive metadata storage.

We recognized three major causes that affect the reusability of publicly available microbiome datasets: 1) **Missing metadata**. A lot of essential metadata are simply not available either because not included among the requested metadata or because not mandatory and hence not compiled by the submitter. One example is the absence of metadata specifying the 16S rRNA hypervariable region amplified and sequenced for most of the studies, which seriously compromise data harmonization efforts. Another information that is often not reported is the presence of an associated publication. The availability of the raw reads on public databases is a requirement for publication in many scientific journals. During the raw reads submission, the researcher is required to provide metadata associated with the dataset, including the presence of a publication. As such, since this step predates the publication itself most of the datasets are uploaded without specifying this information. 2) **Metadata wrongly assigned**. Sometimes metadata can be wrongly assigned to the samples. This can also be the result of mandatory metadata fields that are ambiguous and can lead a researcher inexperienced in the submission process to compile the field in an incorrect way. Wrong metadata can cause the inclusion of wrong datasets into an analysis, potentially affecting the results and leading to incorrect biological conclusions, or, conversely, they can cause the exclusion of datasets from analyses in which they would have fitted. As an example, by comparing the metadata deposited on INSDC with what was reported in the publication we were able to identify studies that wrongly assigned the library strategy as “RNA-Seq” and “WGS” instead of “AMPLICON”.

3) **Inconsistency of the used terminology**. Some metadata fields can be filled with multiple correct metadata leading to inconsistency in the terminology used and affecting the possibility of automatizing the search and filtering of datasets based on these metadata. Good examples are the numerous Taxon ID and scientific names associated with the samples, which are not necessarily wrong, but the lack of consistency in the terms used compromises the usefulness and value of this metadata.

Different works demonstrated the caveats of metadata retrieval and its consequences (*46,47,64*). Researchers have undertaken different approaches to ameliorate this step, in particular using a manual or automated/semi-automated curation (*65*), or developing tools specific for the download of metadata information (*66*). Most of the automated or semi-automated methods are based on Natural Language Processing (NLP) techniques, used to recognize predefined entities in unstructured text, in order to retrieve metadata from the text associated with the samples. Others try to normalize metadata information by grouping or mapping to ontologies (*67–69*). These methods still need a revised step of manual curation and sometimes cannot reconstruct the totality of the metadata associated (*65*). As we demonstrated before, manual curation seems the most accurate solution (*65,70*) if data remains human-readable.

Considering the microbiome field, the INSDC significantly contributed with a recent perspective paper describing the steps that the microbiome research community should take to favor data FAIRification and metadata incorporation (*45*). As microbiome samples are particularly related to the context in which they were collected, data describing measurements or variables related to the context are critical (*45*). Two main subject areas were indicated by the INSDC to improve data standards: i) promote microbiome data sharing and ii) try to remove obstacles and difficulties related to data and metadata submission. Some of their observations and proposals are currently applied by the research community, as for example the “Minimum Information about any (x) Sequence” (MIxS) packages (71) or the incorporation of DOIs for datasets (*72*). Unfortunately, some work is still needed to establish standard procedures and a universal set of ontologies that are easily accessible by the entire community (*45,73*).

In this context, this work also wants to disclose the situation of a sub-field of the microbiome data world: the skin microbiome. The issues revealed by our results show that the search and secondary use of the datasets is still not easy to achieve. Since different studies can rely on different methodologies, different datasets might not be directly comparable and precautions must be taken before combining multiple datasets in a meta-analysis. Without some metadata, a potentially valid dataset can not be included in a meta-analysis. Therefore it is essential for a researcher that wants to valorize a dataset to upload as much information as possible together with the raw reads so as to make the dataset reusable. To motivate researchers in uploading more information, the submission procedure should be made as simple and guided as possible, also to avoid misinterpretations and wrong metadata assignments. To reduce the missingness of metadata, more fields should be made mandatory, such as those referred to the 16S rRNA region sequenced, and new metadata should be included, such as a field that easily discriminates biological samples from negative controls. It also urges the need for standardization of the Taxon ID used in microbiome studies. Guidelines should be given to avoid the use of imprecise Taxon IDs. Efforts should also be made to associate a link to the publication whenever it becomes available, to allow for easier and straightforward access to this resource.

As we have stated, numerous are the aspects related to data and metadata submission that can be improved. Some relate to the submission process itself which can be refined to favor microbiome data reusability, while others strictly depend on the commitment of the researcher performing the submission, who should not overlook the relevance of this step and its importance for the whole scientific community

### The value of a curated skin microbiome collection

Over the past decade, researchers have explored the intricate ecosystem of the skin microbiome (*10*), unveiling the interactions between the microbiome players (bacteria, archaea, fungi and viruses), the skin cells, and the host immune cells that act as barriers, constituting a defense against pathogens invasion and inflammation (*10,74*). Perturbations in the skin ecosystem can cause an unbalance that can even lead to the rising of immune disorders, like allergies, dermatitis or eczema, or chronic injuries, like ulcers. Determining the causes and effects of these processes is not an easy task. Traditional approaches to study skin microbiome mechanisms relies on culture-based techniques, leading to an underestimation of the actors and a bottle-neck selection due to the strict range of cultivable species. The case of *Staphylococcus* genus can serve as an example. Being more easily cultivable than microorganisms belonging to *Corynebacterium* spp. or *Propionibacterium* spp., it would dominate a microbiome dataset, leading to an underestimation of the real biodiversity (*75*). It became obvious that to overcome culture-dependent bottlenecks and to explore the skin microbiome as a whole, a sequencing method must be applied (*10*).

In this context, large-scale sequencing data enable microbiology researchers to obtain deep insights in genetic and functional profiling (*10*) and, nowadays, grand challenges in microbiome science rely on large-scale data science approaches (*20*). Secondary analysis can be full of potential and by-passing the need of generating new large datasets can enormously reduce the costs associated with this kind of study. Impactful meta-analyses have already contributed to advancing the microbiome field, as demonstrated by numerous studies (*17–19*).

From the more applied and clinically relevant studies of skin health and disease to the more theoretical works investigating microbial ecology and the holobiont evolution, all these sub-fields of microbiome research will benefit from the adoption of data-driven approaches based on large-datasets integration (*76*). The availability of a curated collection of microbiome datasets represents the required starting point to make this transition possible and scalable (*23,45*).

Currently, numerous research teams around the world have put efforts in trying to collect and harmonize data from different microbiome fields and various curated collections of microbiome datasets have been published, like the TerrestrialMetagenomeDB (*77*), the HumanMetagenomeDB (*40*), or the Planet Microbe (*78*). Each one of these collections is focused on a specific topic and sometimes on a specific type of data and aims at providing each microbiome research sub-field with a valuable resource to perform data-driven meta-analyses.

Based on these premises and focusing on the skin microbiome sub-field, our work resulted in a comprehensive list of human skin microbiome datasets enriched with metadata information related to the methodological pipelines and the context of the dataset under study.

Skin research produces large quantities of data using a wide range of methods and equipment that require large collaborative efforts. These research endeavors span a broad range of disciplines and are critical to investigating the skin physiology, functions, interactions and health status, from a broad perspective. This can be seen in the bibliometric analysis of published papers related to the datasets retrieved. Research areas and categories from the Web of Science collection and Elsevier’s Scopus classifications showed a scattered distribution of publications in different research areas, but with a higher proportion related to the medicine-related area. As the number of studies grows, it clearly appears that crossing the boundary between medicine and microbial ecology is the lynchpin for a deep understanding of skin health (*4,74*). Indeed, a consistent proportion of the data collected is dedicated to disease conditions, providing valuable material for clinical researchers, but also for microbial ecologists and researchers from other fields of research interested in studying the microbial dynamics in the skin ecological niche. Moreover, taken together, more than half of the studies in our Data Frame 3 collected microbiome data from healthy subjects, providing an invaluable source of information. One of the main challenges for data harmonization is to link the phylogenetic diversity of host-associated microbes to their functional roles within the community and with the host. Much remains to be learned about us as holobionts and much of the information is still kept inside the data.

The curated list we generated can serve as a most comprehensive collection of datasets that can be searched and queried to identify datasets of interest. Researchers interested in conducting meta-analyses with human skin microbiome datasets can use these data frames as a starting point to recover the dataset more suited for their analyses. As demonstrated by the presence of errors in the metadata, these data frames require a curation step. Here, we reported a curated data frame (Data Frame 3) in which we manually corrected errors in the metadata. We also reported two non-curated data frames obtained with the automatic search (Data Frame 1) and with a combination of manual and automatic search (Data Frame 2). These two data frames contain a greater number of studies and samples, however, a careful inspection of these datasets is advised before including any one of those into a meta-analysis.

## Conclusions

The aim of our effort was to help accelerate human skin microbiome research by reducing the amount of time needed to search for datasets and metadata of interest and at the same time favoring data reuse by maximizing the amount of information associated with each dataset. Here we report three data frames containing a comprehensive collection of human skin microbiome datasets enriched with metadata recovered from different sources. The data frames are easily explorable and can be useful for researchers interested in conducting meta-analyses with human skin microbiome amplicon data.

Furthermore, we demonstrated that the reusability of a dataset depends on the amount of information that can be gathered on the dataset itself, that is the amount of metadata deposited by the authors at the time of submission. We are aware that data sharing is increasing throughout the microbiome community, but there are still barriers to making microbiome data truly FAIR. Metadata standards exist, but their proper adoption by the research community is still lagging, as also demonstrated by the NMDC community.

Skin microbiome sampling has the advantage of being non-invasive, easily accessible, and able to provide a huge amount of meaningful information. A curated collection of skin microbiome datasets, enriched with study-related metadata, could be used to investigate health-related phenotypes, offering the potential for non-invasive diagnosis and condition monitoring. Our framework sets the stage for new analyses implementing AI approaches focused on understanding the complex relationships between microbial communities and phenotypes, to predict any condition from microbiome samples. Indeed, considering the skin microbiome topic, a few, very recent works included data integration strategies and AI applications (*79–81*), showing the potential held by these approaches in advancing skin microbiome research.

As the microbiome research field is headed to become a science founded on big-data, the necessity of developing standardized procedures to generate and analyze data acquires importance. The adoption of standard methodologies will help future data integration efforts for the benefit of the whole research community. For this reason, we advocate for a concerted effort to favor standardized microbiome research and exhaustive data sharing.

Further, with this work we want to build a foundation that places microbiome research at the nexus of many subdisciplines within and beyond biology, as for example dermatology, medicine and microbial ecology.

For this reason, this project has the potential to accelerate the development of microbiome-based personalized medicine and non-invasive diagnostics.

## Supplementary data

Supplementary data will be available on our Github repository (https://github.com/giuliaago/SKIOMEMetadataRetrieval).

## Data availability statement

The data frames and supplementaries generated in this study will be available on our Github repository (https://github.com/giuliaago/SKIOMEMetadataRetrieval).

## Author’s notes

G.A. and D.B. equally contributed to this work.

## Authors’ contribution

Conceptualization: G.A., D.B., A.B.; Implementation: G.A., D.B.; Data analysis and visualization: G.A., D.B.; Manuscript preparation: G.A., D.B., A.B.; Review and editing: G.A., D.B., A.B., D.P., M.C., M.L.; Funding acquisition: A.B., M.L.; Supervision: G.A., A.B., M.L.

## Acknowledgements

We thank Intercos Group for their support in developing this manuscript.

